# Triazole Stress Induces Accelerated Degradation of *Aspergillus fumigatus* Hmg1

**DOI:** 10.64898/2026.10.02.756251

**Authors:** Jinhong Xie, Adela Martin-Vicente, Harrison I. Thorn, Uxue Perez-Cuesta, Asmita Nandi, Jarrod R. Fortwendel

**Author notes:** Corresponding Author: Jarrod R. Fortwendel, PhD, University of Tennessee Health Science Center College of Pharmacy, 881 Madison Ave, Rm 338A Memphis, TN 38163.

## Abstract

Triazole antifungals inhibit the lanosterol C14-demethylase Cyp51A/B and are the first-line treatment for invasive aspergillosis caused by *Aspergillus fumigatus*. We previously proposed that triazoles impose a secondary constraint on ergosterol biosynthesis through negative feedback regulation of HMG-CoA reductase (Hmg1) through its sterol sensing domain (SSD), and SSD mutations compromise this regulation to produce resistance. The molecular mechanism underpinning this feedback remained unknown. In this study, we show that voriconazole drives accelerated protein degradation of Hmg1 and that SSD mutation abolishes this regulated degradation. Triazole-induced Hmg1 degradation is proteasome-dependent and partially requires the ERAD E3 ubiquitin ligase, HrdA, and INSIG ortholog, InsA, both conserved regulators of HMG-CoA Reductase abundance. We provide genetic evidence that the initiating negative feedback signal is not triazole acting directly on Hmg1 but lanosterol accumulation resulting from Cyp51A/B inhibition. To test whether depleting Hmg1 protein could re-sensitize resistant strains, we utilized a GFP-targeted degradation system designed to target Hmg1 for degradation without engaging its SSD. Combining this system with voriconazole treatment reduced the MIC at least four-fold in both conidia and established hyphae in susceptible and resistant strains. Together, our findings support a model wherein triazole-induced lanosterol accumulation drives HrdA/InsA-dependent accelerated degradation of Hmg1 and establish that pharmacologically inducing this secondary mechanism can enhance triazole activity in both susceptible and resistant isolates.

## INTRODUCTION

The triazole antifungals are the frontline agents for the treatment of invasive aspergillosis, a life-threatening infection caused predominantly by *A. fumigatus* (1). These drugs inhibit the C14-demethylase Cyp51A/B by binding the heme iron and blocking the oxygen activation required for demethylation, subsequently resulting in disruption of normal sterol homeostasis and accumulation of upstream sterol intermediates (2). In yeasts, the antifungal activity of triazoles is largely fungistatic and depends on Erg3-mediated conversion of accumulated intermediates into a toxic diol (3). In *A. fumigatus*, triazoles are considered fungicidal, yet the molecular basis of this enhanced killing remains incompletely understood. Elsaman *et al*. recently reported that in *A. fumigatus* the fungicidal activity of triazoles correlates with accumulation of eburicol, the Cyp51 substrate, which triggers formation of cell wall carbohydrate patches (4). These findings suggest that the antifungal activity of the triazoles against *A. fumigatus* extends beyond the classical model of ergosterol depletion alone.

Triazole efficacy is increasingly threatened by resistance. Triazole resistance of *A. fumigatus* is consistently associated with higher mortality and has contributed to the designation of *A. fumigatus* as a critical priority pathogen by the World Health Organization (5, 6). The established mechanisms of resistance, including mutations in the *cyp51A* allele and efflux pump overexpression, largely alter the drug-target or intracellular drug accumulation to reduce drug efficacy (7). However, a substantial proportion of resistant clinical isolates carry none of these known resistance determinants (8). Previously, we identified mutations in *hmg1*, encoding HMG-CoA reductase, as a mechanism of clinical triazole resistance not previously described in any fungal pathogen (8). HMG-CoA reductase catalyzes the first committed step of the mevalonate pathway upstream of ergosterol biosynthesis, and the resistance-conferring mutations identified predominantly reside within the predicted sterol sensing domain (SSD) of the protein (8). Since this initial report, *hmg1* SSD mutations have been found in triazole-resistant clinical isolates from multiple countries (9-12).

Although the primary mechanism of action is still deemed to be depletion of ergosterol, we proposed that the triazoles exert a secondary mechanism of action through Hmg1 (13). Inhibition of Cyp51A/B causes accumulation of sterol intermediates that are decoded by the Hmg1 SSD as increased pathway activity, triggering negative feedback regulation of Hmg1. SSD mutations partially disrupt this feedback, resulting in prolonged pathway activity under conditions where it is typically downregulated (13). However, the molecular mechanism through which triazoles impart negative feedback has not been defined. HMG-CoA reductase is regulated by accelerated degradation in both mammalian cells and budding yeast (14), highlighting regulated degradation as a potential mechanism. In contrast, Hmg1 negative feedback in both mammalian and fission yeast systems has also been attributed to regulatory phosphorylation events (15, 16).

In this study, we demonstrate that triazole-induced negative feedback of Hmg1 operates through accelerated protein degradation. Degradation proceeds through the ubiquitin-proteasome system and partially requires the ERAD E3 ligase, HrdA, with the INSIG ortholog, InsA. We provide further evidence that accelerated degradation is likely triggered by lanosterol accumulation induced by Cyp51A/B inhibition rather than through direct triazole-Hmg1 interactions. To explore the translational potential of these findings, we utilized a targeted protein degradation approach to deplete Hmg1 independently of its sterol sensing domains. Targeted Hmg1 degradation reduced the voriconazole MIC by at least four-fold in both conidial and hyphal growth conditions and converted the *hmg1^S305P^* strain from clinically triazole resistant to susceptible. These findings provide proof of concept that targeted degradation of Hmg1 can both enhance triazole activity and restore susceptibility in resistant isolates.

## RESULTS

### Voriconazole accelerates the degradation of Hmg1

We previously proposed that triazoles act through a secondary mechanism by inducing negative feedback regulation of Hmg1 via its SSD (13). In our prior work, we reported that steady-state Hmg1-GFP abundance was not reduced by triazole exposure, which suggested a feedback mechanism operating through targeted protein degradation was not involved. However, steady-state abundance represents the net balance of synthesis and degradation and can therefore remain unchanged even when the rate of degradation is altered. To settle this open question, we measured the impact of triazole stress on Hmg1 degradation directly at the protein level employing cycloheximide (CHX) chase assays. The HiBiT peptide was fused to the C-terminus of the native *hmg1* allele in both the wild-type (*hmg1*-HiBiT) and the S305P SSD mutation (*hmg1^S305P^*-HiBiT) backgrounds. The S305P mutation lies in the middle of the SSD and confers an 8-fold increase in voriconazole MIC (8), which makes this strain a direct test of whether resistance-conferring SSD mutations alter Hmg1 degradation. Both reporter strains retained the triazole-susceptibility profiles of their background strains (Table 1 and Fig. S1), indicating that the HiBiT tag did not perturb Hmg1 function.

**Table 1.** Voriconazole MICs (ug/ml) of strains used in this study.

| Strain | MIC |
| --- | --- |
| *hmg1*- HiBiT | 0.25 |
| *hmg1*<sup>S305P</sup> - HiBiT | 2 |
| Δ*hrdA*/*hmg1*- HiBiT | 1 |
| Δ*insA*/*hmg1*- HiBiT | 0.5 |
| *OE-insA*/*hmg1*- HiBiT | 0.0625 |

Changes in Hmg1 abundance over time were then measured in the presence or absence of voriconazole. Voriconazole was applied at 0.25 × wild-type MIC, a concentration that only slightly inhibited growth, from the start of a 16-h culture. CHX was subsequently added to arrest translation, and mycelia were collected at 0, 2 and 4 h. Relative protein levels were quantitated as the ratio of Hmg1 protein at 2 h or 4 h after CHX addition to the steady-state level (0 h) of the same strain and condition. In the absence of voriconazole, the wild-type strain and *hmg1^S305P^*mutant underwent basal degradation following translational arrest (Fig. 1C). Under voriconazole, the steady-state level (0 h) of wild-type Hmg1 was lower than that of untreated control (Fig. 1A) and the relative level of Hmg1 was significantly lower than that of the untreated control at 4 h (Figs. 1C and 1D). In contrast, although Hmg1^S305P^ presented a higher steady-state abundance under voriconazole, the relative level at 4 h was indistinguishable between the voriconazole-treated and untreated conditions (Figs. 1D), indicating that voriconazole did not accelerate degradation of the mutant protein. Taken together, these results show that voriconazole accelerates the degradation of Hmg1, and that this accelerated degradation is largely abolished by the resistance-conferring SSD mutation S305P. Based on the negative feedback model we proposed previously (13), these findings imply that the feedback regulation of Hmg1 occurs through accelerated protein degradation, and SSD mutation S305P confers resistance by impairing this degradation to escape triazole-induced feedback.

**Figure 1.**
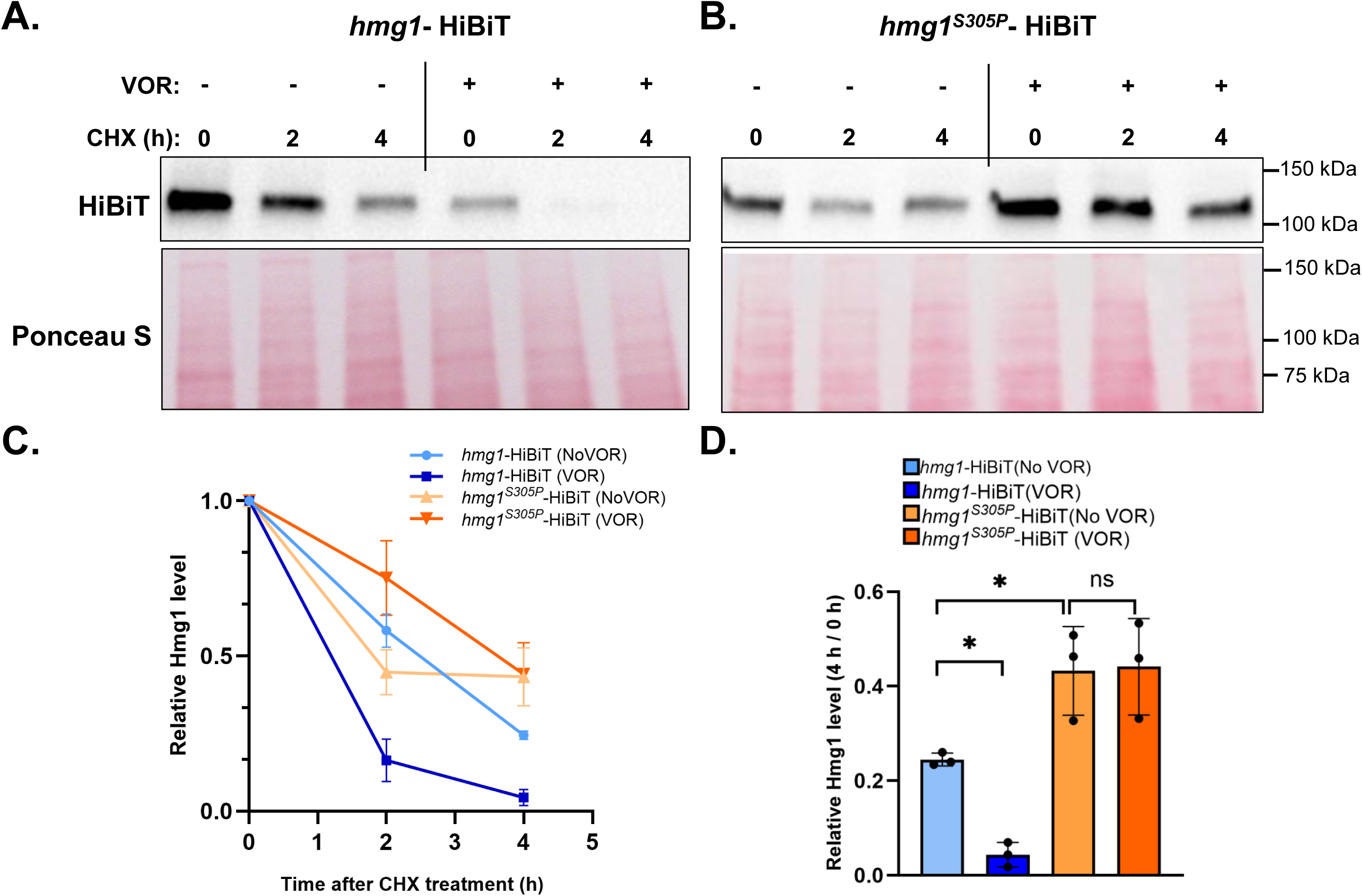
Voriconazole accelerates Hmg1 degradation but not in the *hmg1*^S305P^ mutant. **(A) (B)** Cycloheximide (CHX) chase analysis of Hmg1 degradation in the *hmg1*-HiBiT **(A)** and *hmg1*^S305P^-HiBiT **(B)** strains. Strains were grown for 16 h with 0.0625 µg/ml voriconazole (VOR) or untreated (No VOR). CHX (300 µg/ml) was then added, and mycelia were collected at 0, 2 and 4 h. Hmg1 levels were detected with the Nano-Glo HiBiT Blotting System, and Ponceau S staining was shown as a loading control. **(C)** Relative Hmg1 protein levels over the chase time course from (A) and (B). Hmg1 levels were normalized to Ponceau S and expressed relative to the 0 h time point (steady state) of the same strain and treatment condition. **(D)** Hmg1 protein remaining at 4 h relative to 0 h time point (steady state) for each strain and condition in (A) and (B). Data are mean ± SD of three biological replicates. Statistical significance was assessed by two-way ANOVA. The genotype and VOR treatment interaction was significant (*p* <0.05). Analyses were performed in GraphPad Prism (11.0.2). ∗*p* <0.05, only relevant comparisons are shown.

### Hmg1 accelerated degradation is partially mediated by the ERAD ubiquitin ligase, HrdA

Having established that voriconazole accelerates Hmg1 degradation, we sought to determine the mechanism underlying Hmg1 degradation. In both mammalian cells and *S. cerevisiae*, HMG-CoA reductase is degraded through ER-associated degradation (ERAD), which relies on the ubiquitin-proteasome system (17, 18). Therefore, we reasoned that *A. fumigatus* Hmg1 is also degraded by the ubiquitin-proteasome system. To test this, CHX chase was performed in the presence of MG-132, an inhibitor of the 26S proteasome. MG-132 was added one hour before CHX to ensure proteasome inhibition before protein abundance was tracked. In the presence of voriconazole alone, Hmg1 degraded rapidly and was nearly undetectable by 4 h (Fig. 2A and 2C), as we previously noted. However, co-treatment with MG-132 increased the steady-state level of Hmg1 protein (0 h timepoint) and inhibited Hmg1 degradation over 4 hours (Fig. 2A and 2C). These results show that voriconazole-accelerated degradation of Hmg1 is proteasome-dependent.

**Figure 2.**
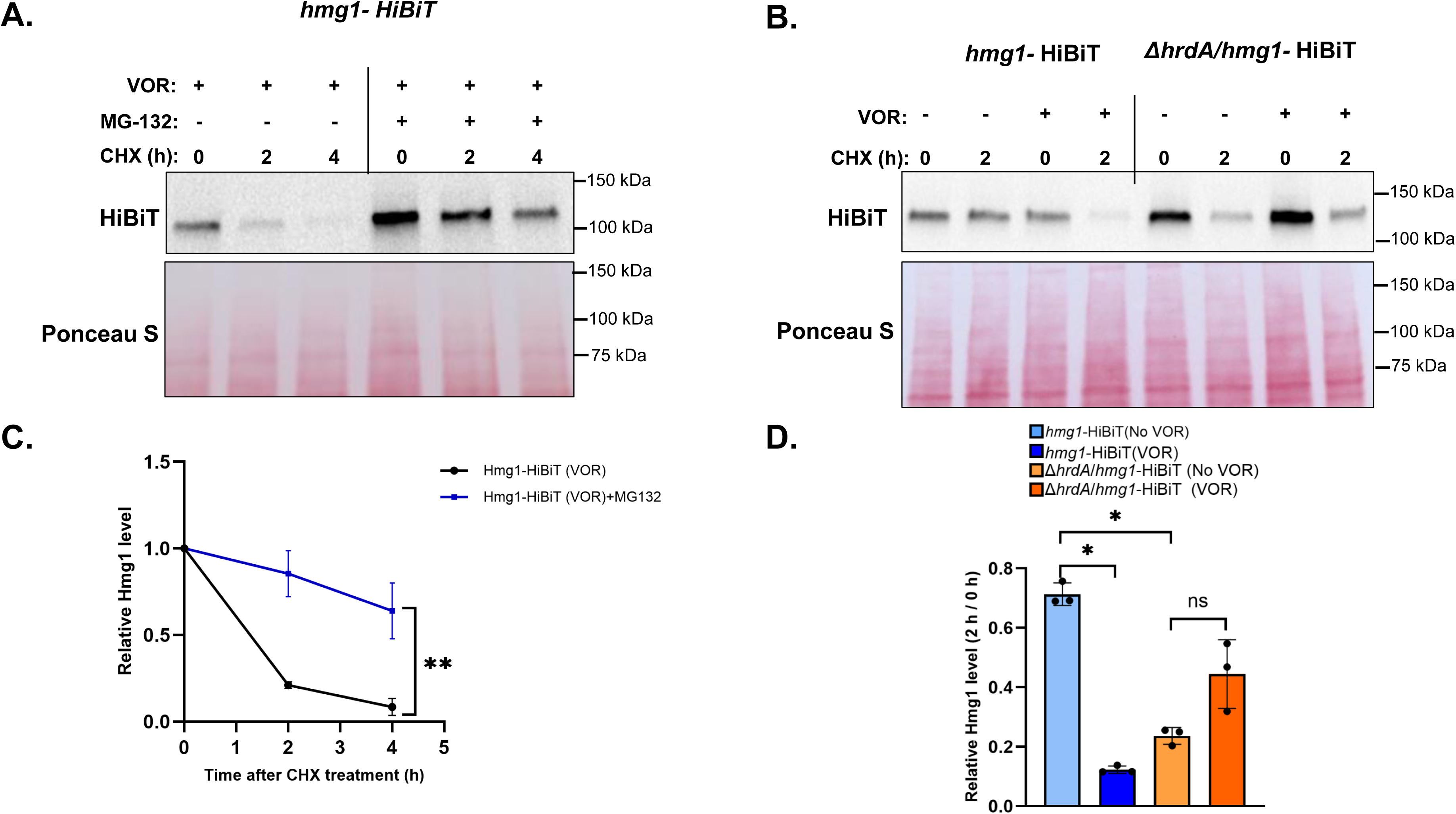
Hmg1 degradation proceeds through the ubiquitin-proteasome system and is partially dependent on HrdA. **(A)** CHX chase analysis of Hmg1 degradation in the hmg1-HiBiT strain in the presence of voriconazole (VOR, 0.0625 µg/ml). Strains were grown for 16 h with voriconazole. MG-132 was added to 40 µg/ml 1 h before CHX (300 µg/ml) and supplemented every 2 h at 40 µg/ml. Mycelia were harvested at 0, 2 and 4 h after CHX addition. **(B)** CHX chase analysis comparing Hmg1 degradation in the *hmg1*-HiBiT and Δ*hrdA*/*hmg1*-HiBiT strains. Strains were grown for 16 h with or without voriconazole (0.0625 µg/ml). CHX was then added, and mycelia were harvested at 0 and 2 h. For all blots, Hmg1-HiBiT was detected with the Nano-Glo HiBiT Blotting System, and Ponceau S staining is shown as loading control. **(C)** Relative Hmg1 protein levels over the chase time course from (A). Hmg1 levels were normalized to Ponceau S and expressed relative to the 0 h time point (steady state) of the same strain and treatment condition. Data are mean ± SD of three biological replicates. Statistical comparisons were performed by two-way ANOVA in GraphPad Prism (11.0.2). The time and MG-132 treatment interaction was significant (*p* < 0.01). **(D)** Hmg1 protein remaining at 2 h relative to 0 h time point (steady state) for each strain and condition in (A) and (B). Data are mean ± SD of three biological replicates. Statistical significance was assessed by two-way ANOVA. The genotype and VOR treatment interaction was significant (*p* < 0.01). Analyses were performed in GraphPad Prism (11.0.2). ∗*p* <0.05, ∗∗*p* < 0.01, only relevant comparisons are shown.

Hrd1 is the E3 ubiquitin ligase that ubiquitinates HMG-CoA reductase in *S. cerevisiae*. It was identified in a screen for *S. cerevisiae* mutants unable to degrade HMG-CoA reductase, taking its name from that phenotype (19). Its mammalian ortholog, *gp78*, carries out the equivalent reaction (20). Further, deletion of *A. fumigatus* ortholog, *hrdA*, has been reported to increase triazole MIC (13, 21). We reasoned that HrdA might serve as the major E3 ligase for Hmg1 degradation (21). To test this, *hrdA* was deleted in the *hmg1*-HiBiT background and Hmg1 degradation was compared between the wild-type and *hrdA*-deletion strains by CHX chase in the presence or absence of voriconazole (Figs. 2B and 2D). Relative levels of protein were quantitated using the Hmg1 protein level after 2 h of CHX treatment normalized to the 0 h steady-state level of the same strain and treatment. Loss of HrdA caused an increase in steady-state levels (0 h timepoint) of Hmg1, suggesting a role for the ERAD system in controlling Hmg1 protein abundance under normal conditions (Fig 2B). In the absence of voriconazole, Hmg1 decreased after translational arrest in both strains, with greater loss observed in the Δ*hrdA* strain by 2 h. Voriconazole further accelerated Hmg1 degradation in the wild type, whereas *hrdA* deletion attenuated the voriconazole-induced Hmg1 degradation (Figs. 2B and 2D). These findings indicate that HrdA contributes to the voriconazole-induced degradation as an E3 ligase. However, Hmg1 continued to decrease following CHX treatment in the Δ*hrdA* strain, implying additional HrdA-independent mechanisms contribute to Hmg1 turnover and may help explain why deletion of *hrdA* increased the voriconazole MIC only from 0.25 to 1 μg/mL, rather than reaching the MIC of 2 μg/mL in *hmg1^S305P^* mutant.

### InsA contributes to the accelerated Hmg1 degradation

The mammalian and budding yeast INSIG orthologs both regulate HMG-CoA reductase turnover in a sterol-dependent manner yet do so in opposite directions. When the sterol signal is high, mammalian INSIG promotes HMGCR ubiquitination by recruiting the E3 ubiquitin ligase gp78, facilitating its degradation (18, 20, 22). In contrast, in budding yeast, INSIG stabilizes the enzyme in response to an accumulated lanosterol signal (23). We previously reported that the *A. fumigatus* ortholog, InsA, may be involved in negative regulation of Hmg1 as its overexpression caused voriconazole hyper-susceptibility (13). How InsA exerts this effect was not mechanistically determined. Given that our data indicates triazoles downregulate Hmg1 through accelerated degradation, InsA was a candidate contributing to this process. To test this, Hmg1 degradation was compared across the wild-type, Δ*insA*, and *insA*-overexpression (OE-*insA*) strains with and without voriconazole by CHX chase assays. For the wild-type and Δ*insA* strains, relative Hmg1 degradation levels were again expressed as the fraction of Hmg1 remaining at 2 h after CHX addition relative to the 0 h time point of the same strain and treatment. As shown in Fig. 3A and 3C, voriconazole significantly lowered the Hmg1 level after 2-hour CHX treatment, again reproducing the accelerated degradation shown in Figure 1. However, no significant difference in Hmg1 degradation was detected between voriconazole treated and non-treated samples in the Δ*insA* background. Notably, the Δ*insA* mutant only displayed a slight increase in voriconazole resistance and did not reach the MIC levels of the *hmg1^S305P^* mutant (Table 1 and Fig.S1). Taken together, these results support a model wherein *insA* deletion does not fully block Hmg1 degradation but reduces the accelerated degradation imposed by voriconazole stress. This is similar to our findings with the Δ*hrdA* mutant, supporting a cooperative mechanism as described in other organisms.

**Figure 3.**
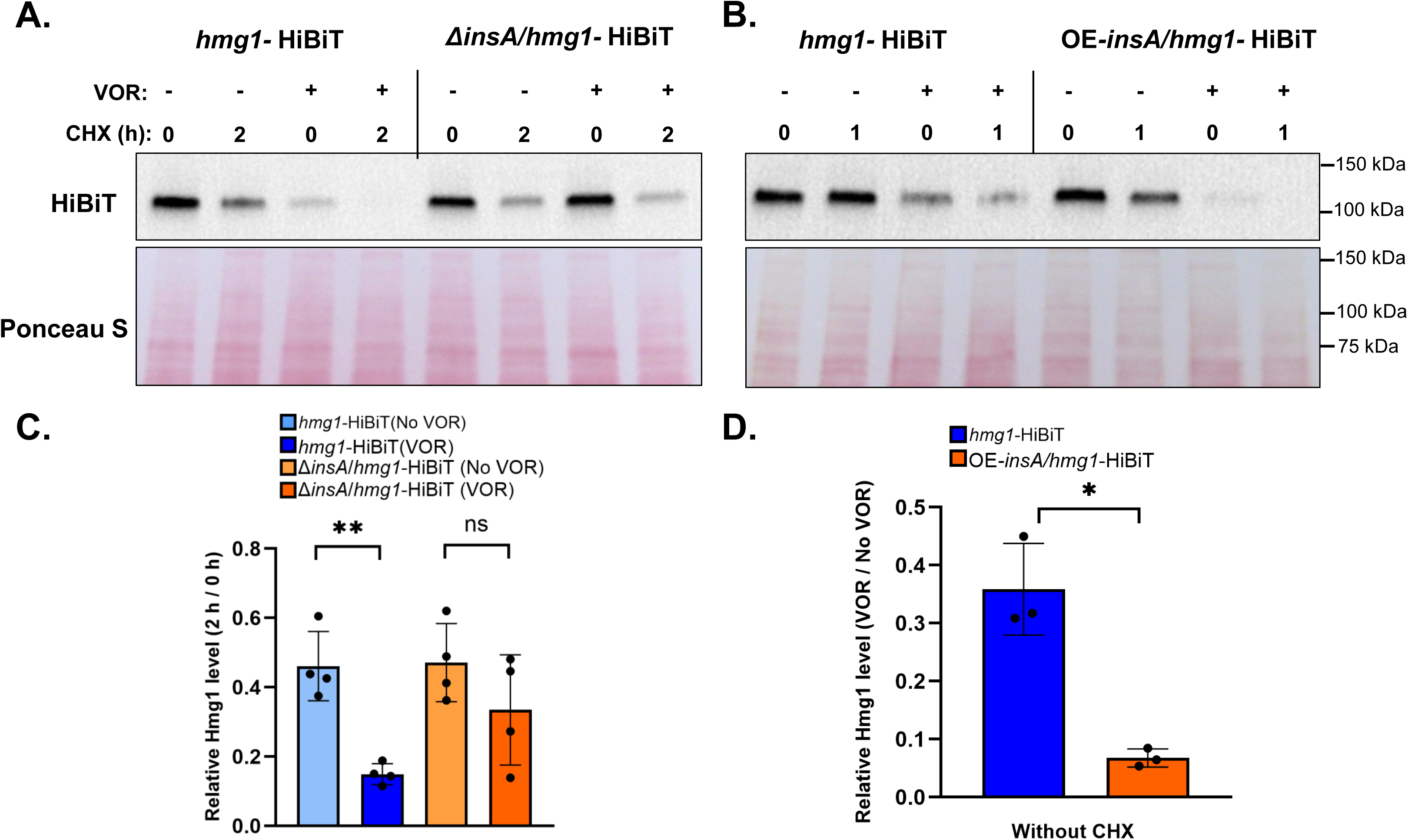
InsA is involved in voriconazole-accelerated Hmg1 degradation. **(A)** CHX chase analysis of Hmg1 degradation in the *hmg1*-HiBiT and Δ*insA*/*hmg1*-HiBiT strains. Strains were grown for 16 h with or without voriconazole (0.0625 µg/ml). CHX (300 µg/ml) was then added, and mycelia were harvested at 0 and 2 h. **(B)** CHX chase analysis of Hmg1 degradation in the *hmg1*-HiBiT and OE-*insA*/*hmg1*-HiBiT strains. Strains were grown for 16 h with or without voriconazole (0.0156 µg/ml). CHX was then added, and mycelia were harvested at 0 and 1 h. **(C)** Hmg1 protein remaining at 2 h relative to 0 h (steady state) for each strain and condition in (A). Data are mean ± SD of four biological replicates. Statistical significance was assessed by two-way ANOVA. The genotype and VOR treatment interaction did not reach statistical significance (*p* = 0.1330). **(D)** Steady-state Hmg1 level (0 h time point) under voriconazole expressed as a fraction of the untreated level for the same strain in (B). Data are mean ± SD of three biological replicates. Statistical comparisons were performed by paired two-tailed *t* test. ∗*p* <0.05, ∗∗*p* < 0.01, only relevant comparisons are shown.

Assuming degradation rate may be increased in OE-*insA*, impacts of *insA* overexpression were quantitated by Hmg1 protein abundance after a shortened 1 h of CHX treatment. However, steady-state levels (0 h) of Hmg1 were barely detectable under voriconazole stress in the OE-*insA* strain (Fig 3B) making contributions of *insA* overexpression to Hmg1 accelerated degradation ambiguous. We instead normalized the steady-state level (0 h time point) of Hmg1 under voriconazole relative to the untreated group, such that a lower ratio reflected greater depletion of Hmg1 by voriconazole. *insA* overexpression sharply reduced this ratio compared to the wild-type background (Fig. 3B and 3D), indicating that increasing InsA promoted Hmg1 depletion under voriconazole. Together, these results suggest that InsA is a component of the pathway that couples triazole stress to Hmg1 accelerated degradation, similar to the role of mammalian INSIG ortholog in the sterol-regulated control of HMG-CoA reductase. We also investigated whether InsA is associated with Hmg1 directly. InsA and Hmg1 interactions were detected through reciprocal co-immunoprecipitation analyses in both the wild-type and S305P backgrounds, and this interaction was detected regardless of voriconazole treatment (Fig. S2). However, whether the S305P mutation or voriconazole treatment alters the strength of the InsA-Hmg1 interaction remains unresolved.

### Triazole-induced lanosterol accumulation is the likely trigger for Hmg1 degradation

Our results so far support the hypothesis that triazoles drive accelerated degradation of Hmg1 through the ubiquitin-proteasome system, but the signal initiating this process remains undefined. Triazoles could promote Hmg1 degradation either directly through a direct drug action on Hmg1, or indirectly by inhibiting the triazole target, Cyp51A/B, and causing the accumulation of sterol intermediates, as we proposed previously (13). To differentiate these two possibilities, we employed mutants expressing the *cyp51A^G54W^* allele. The *cyp51A* G54W substitution resides within the ligand access channel of Cyp51A and sterically hinders the entry of the long-tailed triazoles, such as itraconazole and posaconazole, thereby impairing their binding at the active site of Cyp51A. Enzyme sensitivity to the compact triazoles, such as voriconazole and isavuconazole, is retained (24). Therefore, itraconazole was used in place of voriconazole in this experiment. Since itraconazole is present but Cyp51A^G54W^ remains uninhibited, we reasoned that continued Hmg1 degradation in this background would indicate a direct effect of triazole, whereas stabilization of Hmg1 would indicate that degradation requires inhibition of Cyp51A and subsequent sterol intermediate accumulation. CHX chase assays with itraconazole were performed in the wild-type and *cyp51A^G54W^*. In the wild type, itraconazole significantly decreased the relative Hmg1 level by 4 h compared with the untreated control (Fig.4A and D), consistent with the accelerated degradation observed with voriconazole. In contrast, in the *cyp51^G54W^* background, Hmg1 abundance remained elevated following itraconazole treatment (Fig.4B and D). Therefore, Hmg1 degradation is unlikely to arise from a direct action of triazoles on the Hmg1 and instead depends on the downstream consequences of Cyp51A inhibition.

**Figure 4.**
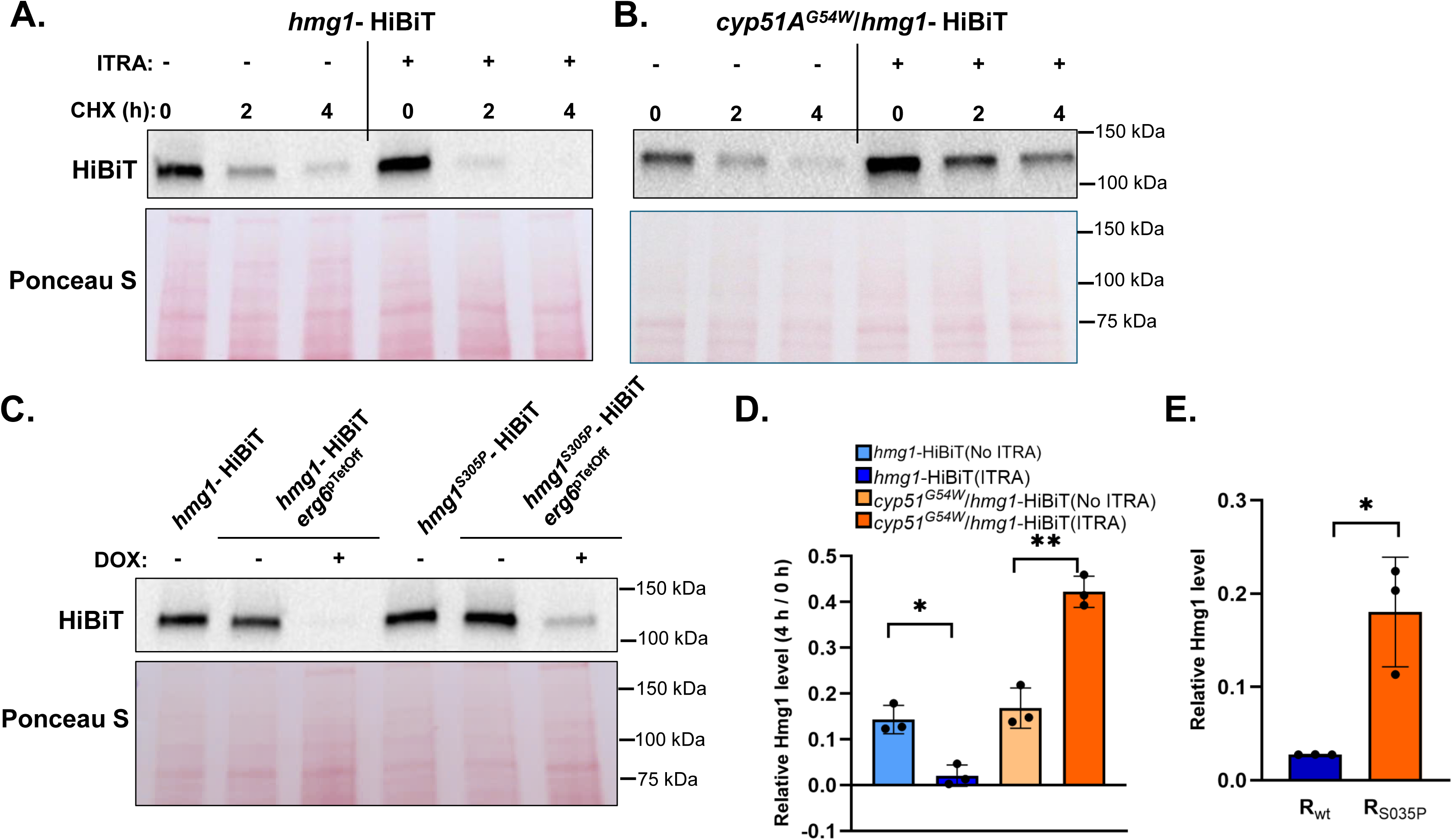
Erg6 repression triggers Hmg1 degradation. **(A) (B)** CHX chase analysis of Hmg1 degradation in the *hmg1*-HiBiT (A) and *cyp51A^G54W^*/*hmg1*-HiBiT (B) strain. Conidia were grown for 16 h with or without itraconazole (ITRA, 0.03125 µg/ml). CHX (300 µg/ml) was then added, and mycelia were harvested at 0, 2 and 4 h. **(C)** Steady-state Hmg1 levels in the *hmg1*-HiBiT/*erg6*^pTetOff^ and *hmg1*^S305P^-HiBiT/*erg6*^pTetOff^ strains cultured with or without doxycycline (DOX, 0.5 µg/ml) to repress *erg6* and induce lanosterol accumulation in the absence of triazole. For all blots, Hmg1-HiBiT was detected with the Nano-Glo HiBiT Blotting System, and Ponceau S staining is shown as loading control. **(D)** Hmg1 protein remaining at 4 h relative to 0 h (steady state) for each condition in (A) and (B). Data are mean ± SD of three biological replicates. Statistical comparisons were performed by two-way ANOVA. The genotype and ITRA treatment interaction was significant (*p* < 0.01). **(E)** Quantification of Hmg1 abundance from (C), normalized to Ponceau S and expressed as the ratio of Hmg1 in the DOX-treated to the untreated condition within each background (R = Hmg1+DOX / Hmg1−DOX). Data are mean ± SD of three biological replicates. Statistical comparisons were performed by paired two-tailed *t* test. Analyses were performed in GraphPad Prism (11.0.2). ∗*p* <0.05, ∗∗*p* < 0.01, only relevant comparisons are shown.

In both mammalian and yeast systems, sterol and / or isoprenoid intermediates have been found to serve as a feedback signal for HMG-CoA reductase, though both the molecules involved and their regulatory functions are different. In mammalian cells, lanosterol and 24,25-dihydrolanosterol are the major signals that accelerate degradation of HMG-CoA reductase (25, 26). In *S. cerevisiae*, geranylgeranyl pyrophosphate (GGPP) accelerates Hmg2 degradation, whereas lanosterol promotes the INSIG ortholog Nsg1 to stabilize the enzyme (23, 27). We previously reported that voriconazole treatment leads to accumulation of lanosterol and eburicol (13). We therefore reasoned that the accumulation of sterol intermediates might be the signal that triggers Hmg1 degradation. Because exogenously supplied sterol intermediates are poorly taken up by *A. fumigatus*, genetic manipulation was used to drive their accumulation. We previously showed that repression of *erg6*, the essential sterol methyltransferase functioning just upstream of Cyp51A, induces the accumulation of lanosterol without altering other intermediates (28). We placed *erg6* under the control of a doxycycline-repressible promoter (*erg6^pTetOff^*) in both the wild-type and *hmg1*^S305P^ backgrounds and measured steady-state Hmg1 abundance. Hmg1 levels were normalized to their untreated counterparts in each background. In the wild-type background, repression of *erg6* alone was sufficient to reduce Hmg1 to nearly undetectable levels in the absence of triazole (Fig. 4C and E). In the *hmg1^S305P^* background, Hmg1 was retained at a significantly higher relative level under the same condition (Fig.4C and E). These results strongly suggest that accumulation of lanosterol is sufficient to lower Hmg1 abundance without triazole exposure, and that the S305P mutation blunts this response.

### Targeted degradation of Hmg1 resensitizes resistant strains to triazoles

Hmg1 catalyzes the first rate-limiting step of the ergosterol biosynthesis pathway, making it an attractive antifungal target. Statins inhibit mammalian HMG-CoA reductase by competing at its catalytic domain and also exhibit antifungal activity against *A. fumigatus*. However, this activity requires concentrations far exceeding those achievable clinically, which limits their usage as antifungals (29). Our results above suggest the usefulness of employing a different route to disabling the enzyme, in which Hmg1 is degraded rather than inhibited, to improve co-therapy with triazole antifungals. We therefore reasoned that forcing depletion of Hmg1 would arrest pathway activity, impair fungal growth, and subvert triazole resistance.

To achieve targeted degradation, we adopted a GFP-directed protein degradation system (30), in which a nanobody fused to the E3 ubiquitin ligase Rnf4 recognizes GFP-tagged proteins and delivers them to the ubiquitin-proteasome machinery. The doxycycline-inducible construct (TetOn-GFPNb-Rnf4) was introduced into the parental strains and into the *hmg1*-GFP and *hmg1^S305P^*-GFP backgrounds, which were previously generated in our laboratory and shown to be functional (13). Doxycycline induction of GFPNb-Rnf4 had no effect on colony growth in the absence of a GFP-tagged target and neither GFP-tagged strain was affected for growth in the absence of the degrader construct (Fig. S3A). In contrast, both *hmg1*-GFP and *hmg1^S305P^*-GFP strains carrying the inducible GFPNb-Rnf4 construct developed pronounced growth defects in the presence of doxycycline, with growth abolished at 10 µg/ml (Fig. S3A).

Hmg1-GFP abundance was then measured under two doxycycline dosing regimens to test effects of expressing the degrader construct from the time of conidial germination and after established hyphal development. To do so, a low concentration of doxycycline (0.3 µg/ml), sufficient to restrict but not abolish colony growth, was added at the start of culture, whereas a high concentration (100 µg/ml) was added after 16 h of initial culture in doxycycline free media, respectively. Steady-state Hmg1-GFP levels decreased under both regimens when compared to uninduced controls, with Hmg1 protein abundance falling to nearly undetectable levels within 2 h of the high-dose treatment (Fig. 5A and B). Significant degradation was therefore achieved in both developing and established hyphae and likely underpins the loss of growth seen in agar-based growth assays (Fig. S3) The effect of induced Hmg1 depletion on triazole susceptibility was measured by supplying doxycycline during voriconazole MIC testing under the same two dosing regimens. In the wild-type *hmg1*-GFP strain, the MIC was reduced from 0.25 to ≤0.0625 µg/ml in both conditions (Table 2). In the resistant *hmg1^S305P^*-GFP strain, the MIC was reduced from 2 µg/ml to 0.5 µg/ml when doxycycline was supplied from the start of culture, and to between 0.25 and 0.5 µg/ml when it was supplied after 16 h. Thus, degradation of Hmg1 lowered the MIC of the resistant strain below the CLSI breakpoint for voriconazole, suggesting that induction of Hmg1 degradation through a mechanism that subverts the intrinsic machinery regulating this process is sufficient to overcome the resistance conferred by SSD mutation.

**Figure 5.**
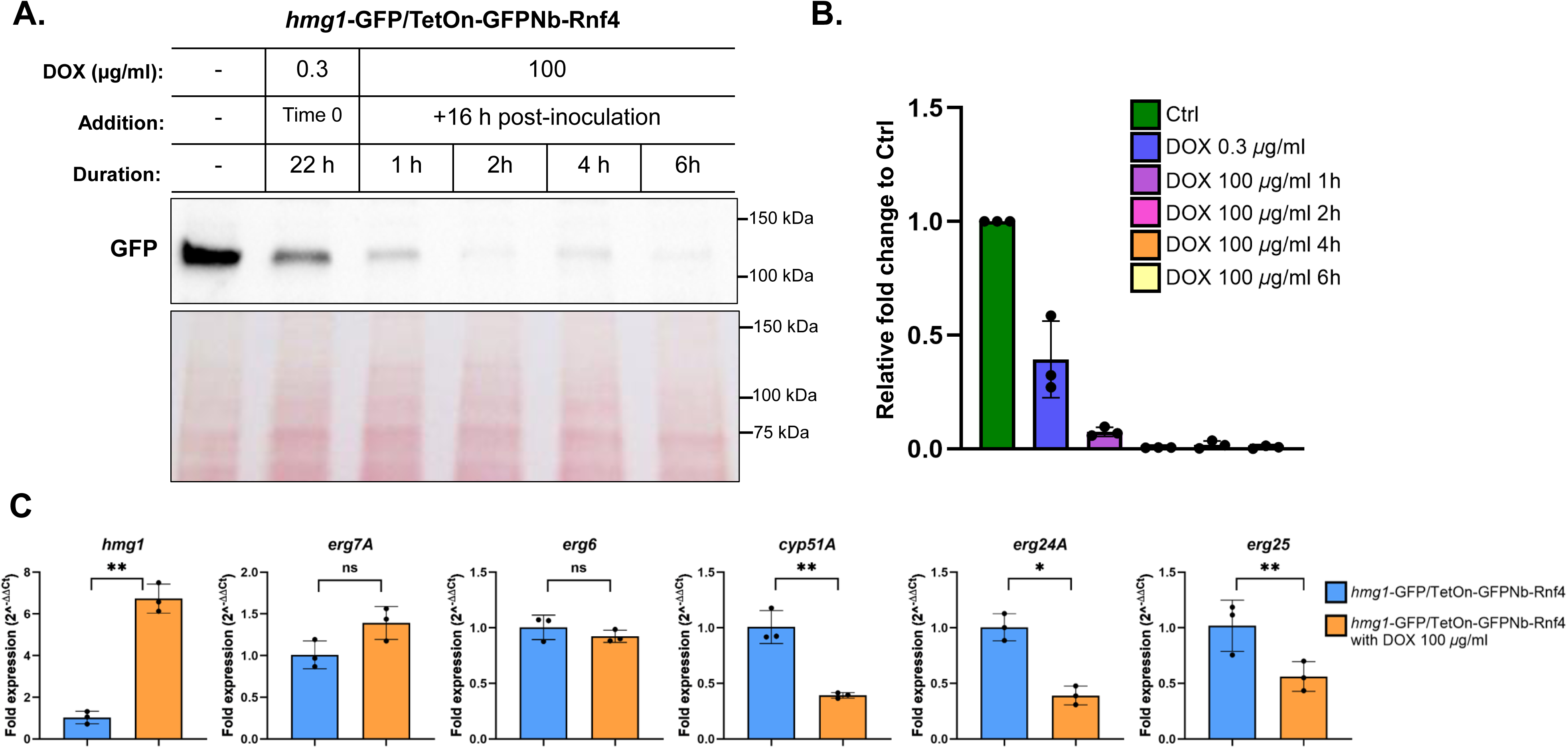
Targeted degradation of Hmg1 depletes the protein irrespective of growth phase and induces compensatory *hmg1* transcription. **(A)** Hmg1-GFP protein levels in the *hmg1*-GFP/TetOn-GFPNb-Rnf4 strain under two doxycycline additions. For the low-dose regimen, doxycycline (DOX, 0.3 µg/ml) was added at the start of culture and maintained for 22 h. For the high-dose regimen, conidia were grown for 16 h without doxycycline to develop hyphae, after which doxycycline (100 µg/ml) was added and mycelia were harvested at 1, 2, 4 and 6 h. An untreated control (no DOX) was grown for the same total duration. Hmg1-GFP was detected by anti-GFP immunoblotting, and Ponceau S staining is shown as a loading control. **(B)** Quantification of Hmg1-GFP band intensities from (A), normalized to Ponceau S and expressed relative to the untreated control. **(C)** RT-qPCR analysis of *hmg1*, *cyp51A*, *erg7A*, *erg6*, *erg24A* and *erg25* transcript levels in the *hmg1*-GFP/TetOn-GFPNb-Rnf4 strain. Hyphae were grown for 16 h without doxycycline, then cultured for an additional 4 h with or without doxycycline (100 µg/ml). Relative expression was determined by the 2^−ΔΔCt^ method using *tubA* as the reference gene. Data are mean ± SD of three biological replicates. Statistical significance was determined by comparing ΔCt values using a paired *t*-test. ∗*p* <0.05, ∗∗*p* < 0.01, only relevant comparisons are shown.

**Table 2.**
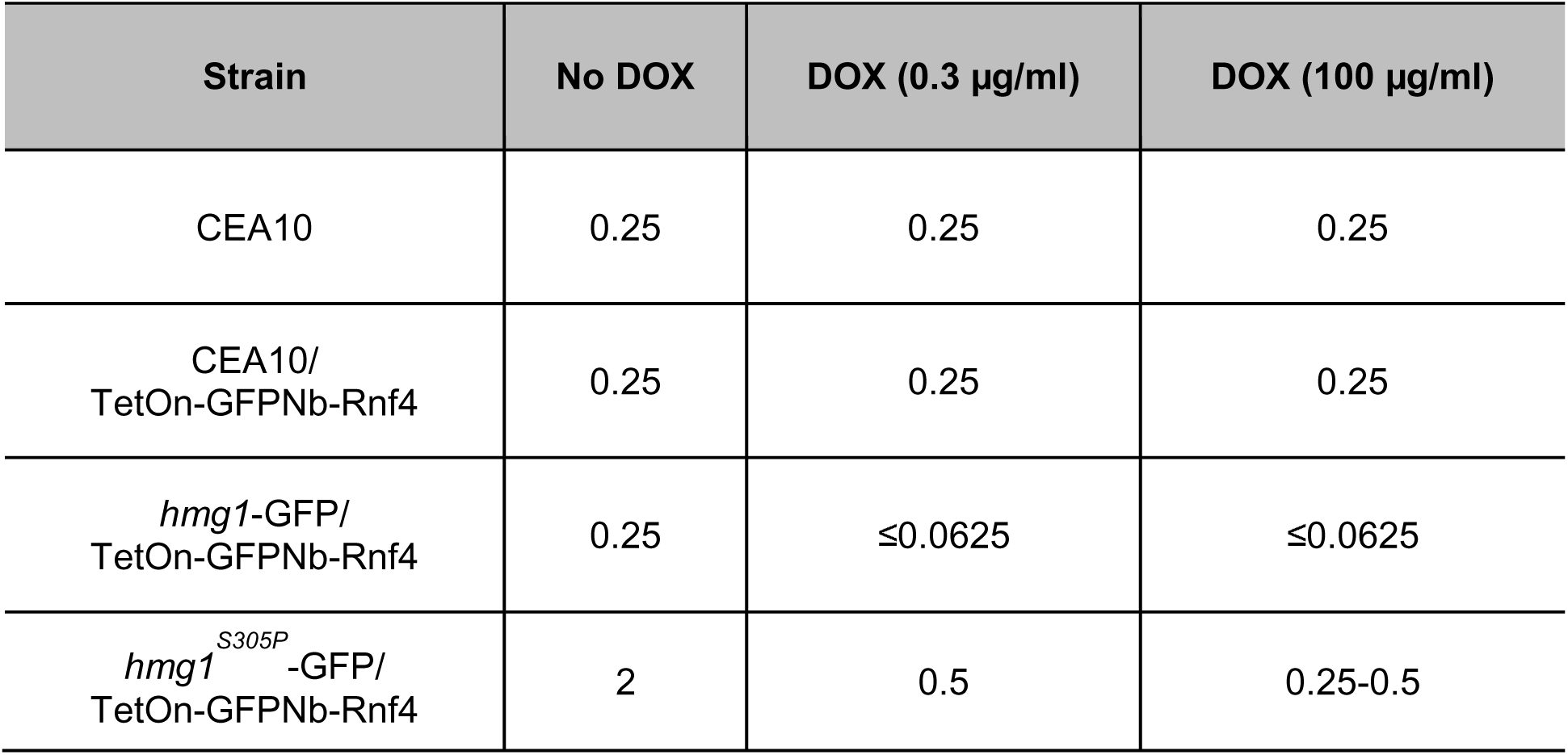
Voriconazole MICs (ug/ml) after targeted degradation of Hmg1.

To examine how loss of Hmg1 affects the ergosterol biosynthesis pathway, we induced its degradation and measured the resulting expression changes of pathway genes. Doxycycline was added to 16-h hyphae of the *hmg1*-GFP/TetOn-GFPNb-Rnf4 strain for 4 h, and transcripts of *hmg1* together with downstream pathway genes were quantified by RT-qPCR. Degradation of Hmg1 significantly increased the *hmg1* transcript, whereas *cyp51A*, *erg24A* and *erg25* were significantly decreased, and *erg7A* and *erg6* were unchanged (Fig. 5C). Taken together, these results show that degrading Hmg1 selectively downregulates *cyp51A* and downstream ergosterol biosynthesis genes and is accompanied by increased *hmg1* transcription, implying that the cell compensates at the transcriptional level while the protein is depleted.

## DISCUSSION

The triazole antifungals inhibit Cyp51A/B and our previous work proposed that this inhibition places a second constraint on ergosterol biosynthesis by inducing negative feedback regulation of Hmg1 (13). That model rested on genetic and phenotypic evidence and left the mechanism of the feedback undefined. Accelerated degradation is a conserved mode of HMG-CoA reductase regulation in both mammalian cells and budding yeast (17, 18), which made degradation a candidate mechanism. We previously reported steady-state Hmg1 abundance using GFP levels as a proxy and found no decrease under sublethal voriconazole stress (13). However, the readout might be masked by the free GFP released during Hmg1-GFP degradation. Moreover, steady-state abundance is the net balance of synthesis and degradation, and a rise in synthesis can offset an increase in degradation resulting in no change of the amount of protein present. In support of this, we find here that forced degradation of Hmg1 for a prolonged time course significantly induced *hmg1* transcript (Fig. 5C), showing that *A. fumigatus* responds to loss of the protein by increasing its expression.

The data collected in this study further support and clarify our model for Hmg1 feedback (Fig.6). Following translational arrest, voriconazole accelerates the degradation of Hmg1 in the wild type relative to the untreated control (Fig.1), which indicates that the negative feedback is mediated at the level of protein degradation. The initiating signal is likely not the drug itself but sterol intermediates, particularly lanosterol, that accumulate once Cyp51A/B is inhibited (Fig.4C). InsA is at least partially required for this acceleration, and increased InsA dramatically promotes depletion of Hmg1 under voriconazole (Fig.3). The accelerated degradation is proteasome dependent and requires HrdA, a predicted E3 ligase ortholog, although the incomplete block in the Δ*hrdA* mutant indicates that it is not the only component required (Fig.2B). In the *hmg1^S305P^*mutant, physical association between InsA and Hmg1 remains detectable, yet voriconazole-induced degradation is blunted, so the pathway activity is maintained and the strain remains resistant.

**Figure 6.**
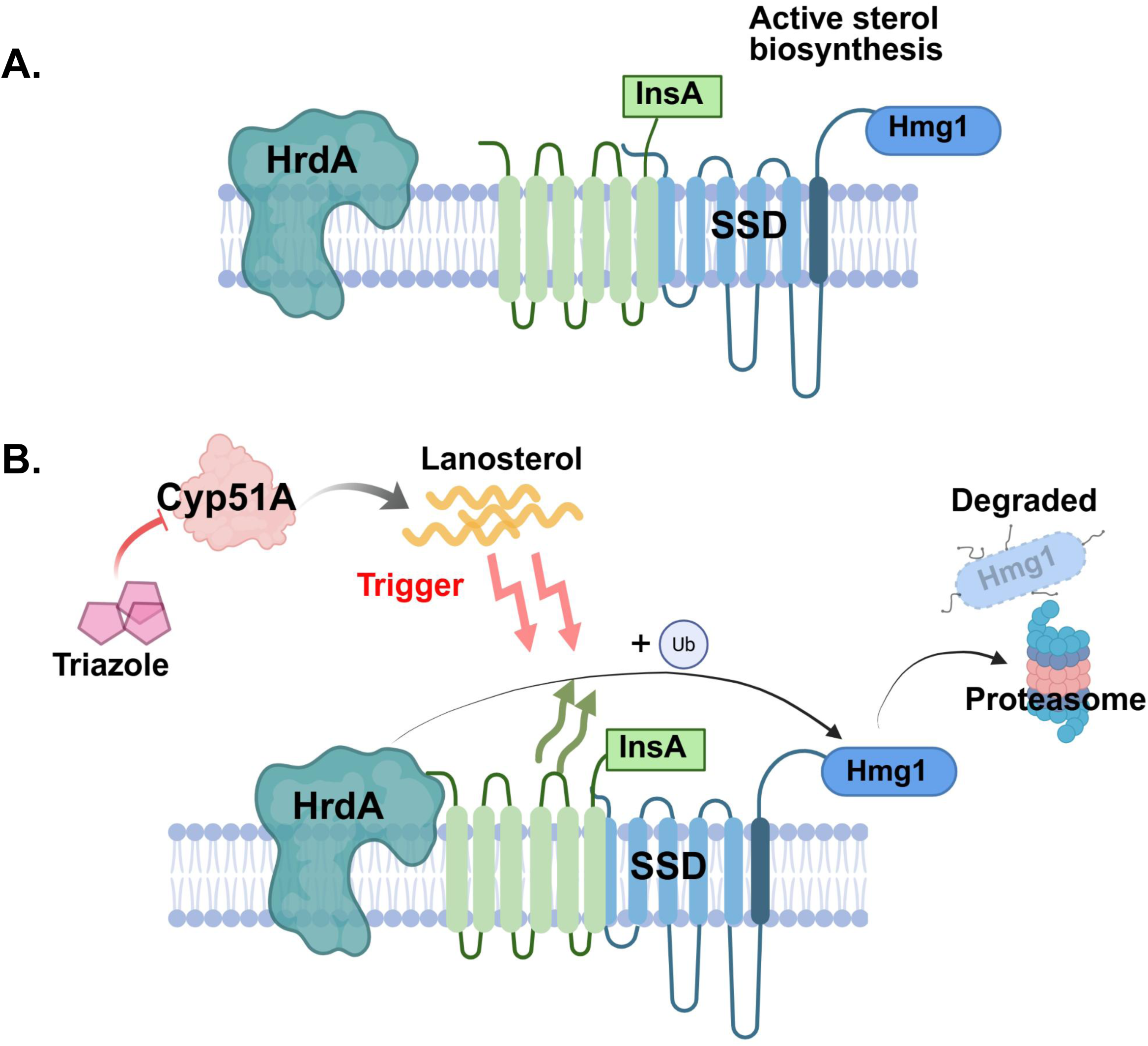
Proposed model for triazole-induced accelerated degradation of Hmg1. **(A)** Under basal conditions, Hmg1 resides in the ER membrane with InsA associated. Sterol biosynthesis proceeds normally and Hmg1 remains stable. **(B)** Triazole inhibition of Cyp51A blocks lanosterol demethylation and causes lanosterol accumulation. This accumulation is accompanied by InsA-dependent, HrdA-mediated Hmg1 degradation through proteasome, resulting in downregulation of sterol biosynthesis. The S305P substitution in the SSD does not disrupt the association between InsA and Hmg1 but blocks the accelerated degradation, allowing the mutant protein to persist under triazole stress and conferring resistance.

To begin delineating the relationship of putative regulatory components, we previously deleted *hrdA* in an *insA* overexpression background and observed no change in voriconazole susceptibility compared to the *insA*-overexpressing strain (13). That comparison relied on a susceptibility readout, which reports the consequence of genetic manipulation and cannot isolate the contribution of a single ligase. In addition, the earlier measurement was made in a strain expressing *insA* more than fifty-fold above its normal level (13), which may mask a partial contribution of HrdA to the response. In this study, we tested roles for each of these conserved HMG-CoA Reductase regulators by measuring Hmg1 protein level directly. Voriconazole accelerates Hmg1 degradation in the wild type, but this was blunted in the Δ*hrdA* strain (Fig. 2B and 2D), indicating HrdA participates in the voriconazole-induced degradation of Hmg1. However, deletion of *hrdA* raised the voriconazole MIC four-fold relative to the wild type, whereas the *hmg1^S305P^* mutation conferred an eight-fold increase (Table 1 and Fig. S1). This difference suggests that degradation is reduced rather than fully abolished when the ligase is absent. This HrdA-independent degradation could help explain why deletion of *hrdA* failed to reverse the voriconazole hypersusceptibility caused by *insA* overexpression in our previous study. Several ubiquitin ligases, such as gp78, Trc8, and RNF145 act on HMG-CoA reductase in mammalian cells, and the loss of any one of them only partially impacts degradation (31, 32). Therefore, HrdA is likely to be one of the E3 ligases acting on Hmg1 and additional E3 ligases may also contribute to its degradation.

Our findings also support the hypothesis that accelerated degradation of Hmg1 is triggered not by the triazoles acting on Hmg1 directly, but by the consequences of their inhibition of Cyp51. When the *cyp51A*^G54W^ substitution obstructs the ligand access channel and prevents the long tail triazole itraconazole from inhibiting Cyp51A, itraconazole no longer accelerated Hmg1 degradation (Fig. 4B). However, *erg6* repression, which causes lanosterol accumulation selectively, was sufficient to deplete Hmg1 with no triazole present (Fig. 4C). The S305P substitution blunted this depletion (Fig. 4C and 4E), placing the sterol sensing domain between the lanosterol signal and the loss of the protein. In mammals, lanosterol and 24,25-dihydrolanosterol accelerate INSIG-dependent degradation of HMG-CoA reductase, and the activity has been traced to the pair of methyl groups at C4 of the sterol ring (25, 33). Eburicol accumulates alongside lanosterol under triazole stress and carries the same C4 dimethyl group, so it may contribute to the response as well, though this was not tested here. The budding yeast system reads a different molecule for an effect in the opposite direction. There, GGPP drives Hmg2 degradation through the HRD pathway, while lanosterol instead stabilizes the enzyme by promoting its association with the INSIG ortholog, Nsg1 (23, 27). Thus, our findings parallel the mammalian arrangement, in which both lanosterol and InsA promote degradation. Lanosterol may serve as a feedback signal because it is the first sterol-type intermediate of the pathway, and it lies upstream of the oxygen-intensive demethylation catalyzed by Cyp51 (34). In mammals, nine of the eleven molecules of O_2_ consumed during the synthesis of cholesterol are spent on the demethylation reactions downstream of lanosterol (34). Hypoxia impairs lanosterol demethylation and induces Insig through hypoxia-inducible factor (HIF)-1α to accelerate HMG-CoA reductase degradation (35)*. A. fumigatus* is an obligate aerobe, and ergosterol synthesis requires multiple oxygen-dependent reactions downstream of lanosterol. Feedback at lanosterol would allow the cell to slow down sterol synthesis before committing oxygen to these reactions. Consistent with this possibility, artificially induced Hmg1 degradation selectively reduced the transcription of *cyp51A* and downstream ergosterol biosynthetic genes *erg24A* and *erg25* (Fig.5C). Together, these observations suggest that lanosterol-mediated Hmg1 degradation may help coordinate sterol biosynthesis with cellular oxygen utilization.

Under voriconazole, Hmg1 remained relatively stable when *insA* was deleted and decreased to near-undetectable levels when *insA* was overexpressed (Fig. 3A and 3B). These findings indicate that InsA participates in triazole-induced Hmg1 degradation, but degradation still proceeds in the absence of InsA, suggesting that an InsA-independent pathway also operates. Both mammalian cells and *S. cerevisiae* sense two signals to regulate HMG-CoA reductase degradation. In mammalian cells, lanosterol and 24,25-dihydrolanosterol initiate SSD/INSIG-dependent ubiquitination of HMGCR, but ubiquitination alone is not sufficient for degradation (34). Sterols also induce UbiA prenyltransferase domain-containing protein 1 (UBIAD1) binding to HMGCR in the ER and blocking degradation. Accumulated GGPP releases UBIAD1 from HMGCR, which allows HMGCR to undergo membrane extraction and degradation (36, 37). In *S. cerevisiae*, the two signals oppose one another. GGPP induces Hmg2 ubiquitination and degradation, whereas lanosterol promotes association of Nsg1 with Hmg2 and stabilizes it (23). Whether *A. fumigatus* also integrates a second signal, and whether this signal acts in the same direction as InsA or opposes it remains unknown. The exact biological function of SSD remains undetermined. In both systems an intact SSD is required for the signal-induced conformational change that precedes ubiquitination and degradation (38, 39). In *A. fumigatus*, it could be needed to sense the sterol signal, to convert that signal into a conformational change, or to permit ubiquitination once InsA is bound. Further study is needed to dissect the whole process. Most established mechanisms of triazole resistance operate through either changes to the drug targets, such as mutations in *cyp51A* allele, or reduced intracellular drug concentration through efflux pump overexpression (7). The S305P SSD mutation prevents the triazole-induced degradation of Hmg1, removing the additional restriction on sterol pathway and consequently conferring resistance. When *hmg1* SSD and *cyp51A* mutations co-occur, as they frequently do in clinical isolates (13), they each remove a different inhibitory effect of triazoles, and the resulting resistance exceeds that conferred by either mutation alone (11).Similar HMG-CoA reductase SSD-mediated resistance has not been reported in pathogenic yeasts. The regulation of Hmg2 in *S. cerevisiae* may help explain this absence. In *S. cerevisiae*, triazole-induced lanosterol accumulation stabilizes rather than degrades Hmg2 (23). Therefore, SSD mutation could not cause resistance by removing this inhibitory effect.

Our findings also show that using a doxycycline-inducible GFP-targeting protein degradation construct to mimic pharmacological depletion of Hmg1 causes growth arrest and lowered voriconazole MICs. Importantly, this approach reduced triazole MIC in the resistant *hmg1^S305P^* mutant to below the epidemiological cutoff value for susceptibility (Table 2). Therefore, reducing Hmg1 activity or abundance enhances triazole efficacy. Statins compete at the catalytic domain and have been shown to act additively with triazoles against *A. fumigatus*, although the effective statin concentrations remain above clinically achievable levels (29, 40). Degradation-inducing compounds have also been developed in the mammalian system. The synthetic sterol analog Cmpd 81 (also named as HMG499) induces HMGCR degradation (33). However, Cmpd 81 operates through the endogenous Insig-dependent pathway and would be expected to face the same SSD mutations that confer triazole resistance in *A. fumigatus* (41). Targeted protein degradation through a heterologous degradation machinery offers a different pharmacological strategy. Proteolysis-targeting chimeras (PROTACs) are heterobifunctional molecules that bring a target protein into proximity with an E3 ubiquitin ligase, leading to targeted protein degradation, and their application in antimicrobial drug discovery has been proposed as a route to overcome resistance that defeats conventional inhibitors (42). Won et al. demonstrated this principle in mycobacteria, where directing essential proteins to the ClpC1P1P2 protease through induced proximity inhibited growth and potentiated existing antibiotics (43). The genetic degradation system used in this study provides proof of concept that depleting Hmg1 through a pathway independent of its sterol sensing domain can reverse triazole resistance in *A. fumigatus*. This approach could be an efficacious standalone or co-therapeutic antifungal strategy to combat invasive aspergillosis.

## METHODS

### Strains and growth conditions

All genetic manipulations were performed in the *A. fumigatus* parental strain CEA10. All strains used in this study are listed in Table S1. Strains were routinely maintained at 37 °C on glucose minimal medium (GMM) agar as previously described (44). Conidia were harvested from 3-day-old cultures in sterile water, stored at 4 °C, and enumerated using a hemocytometer for use. Unless otherwise indicated, all solid cultures were grown on GMM agar and all liquid cultures were grown in GMM supplemented with 0.5% (w/v) yeast extract (GMM+YE).

### Strain construction

All strains were generated in the CEA10 background using a CRISPR-Cas9 gene editing system, as previously described (45). CRISPR RNAs (crRNAs) targeting protospacer-adjacent motif (PAM) sites at or flanking each locus of interest were designed for each modification. All crRNA target sequences and repair-template primers used for each strain are listed in Table S2. RNP complexes were assembled *in vitro* from synthetic crRNA, tracrRNA and recombinant Cas9 following established protocols. Equimolar crRNA and tracrRNA were combined in duplex buffer and annealed at 95 °C for 5 min, and the resulting gRNA duplex was complexed with Cas9 for 5 min at room temperature.

Repair templates were amplified by PCR with primers incorporating 40-bp microhomology arms matching the 5′ and 3′ ends of the target locus, and the specific modification was determined by the template used. (i) Gene deletions (Δ*hrdA* and Δ*insA*) were generated by targeting two PAM sites flanking the open reading frame and replacing it with a selectable resistance cassette. (ii) Promoter replacements were generated by targeting a PAM site immediately upstream of the start codon and replacing the native promoter with the *hspA* promoter for overexpression (*insA^OE^*) or with the pTetOff promoter for doxycycline-repressible expression (*erg6^pTetOff^*). (iii) C-terminal HiBiT, GFP and 3×FLAG fusions were introduced at the native 3′ end of *hmg1* or *insA* by targeting a PAM site near the stop codon. Repair templates were generated by fusion PCR joining the 3′ UTR homology arm to a resistance cassette, with the in-frame tag encoded in the 5′ extension of the 3′ UTR forward primer. (iv) Point mutations (*hmg1^S305P^*) were introduced using repair templates carrying the desired codon change. (v) The doxycycline-inducible GFPNb-Rnf4 degron cassette (TetOn-GFPNb-Rnf4) was integrated at the *fcyB* locus.

Protoplast generation and transformation were performed as described (45). Briefly, the transformation mixture was assembled on ice by combining 5 μl RNP, 900 ng repair template and 3 μl polyethylene glycol (PEG)-CaCl_2_ buffer with 10 μl protoplasts (1-5 × 10[) in STC buffer (1.2 M sorbitol, 7.55 mM CaCl_2_·H_2_O, 10 mM Tris-HCl, pH 7.5). After 50 min on ice, 57 μl PEG-CaCl_2_ buffer was added and the mixture held at room temperature for 20 min, then brought to 200 μl with STC buffer and spread on SMM agar. Plates were held at room temperature overnight before applying a selective SMM top-agar overlay containing the appropriate drug and incubated at 37 °C until colonies formed. Transformants were single-colony purified and confirmed by PCR across the 5′ and 3′ integration junctions; in-frame tag fusions and point mutations were additionally verified by Sanger sequencing.

### Membrane protein extraction

All protein samples were prepared for membrane fractions. Cultures grown in GMM+YE were harvested by filtration through Miracloth, washed twice with sterile water, and either processed immediately or stored at -80 °C. Mycelia were ground to a fine powder under liquid nitrogen and resuspended in extraction buffer (50 mM HEPES, 50 mM EDTA) containing 1 mM Pefabloc and 1× protease inhibitor cocktail (Sigma-Aldrich, P8215). After 20 min on ice with brief vortexing every 5 min, lysates were cleared at 8,000 r.p.m. for 20 min at 4 °C, and the supernatant was centrifuged at 100,000 × *g* for 1 h at 4 °C to pellet membranes. The membrane pellet was resuspended in 1× RIPA lysis buffer (Millipore, 20-188) supplemented with 10% (v/v) glycerol, solubilized by end-over-end rotation for 30 min at 4 °C, and clarified at 8,000 r.p.m. for 2 min at 4 °C. Protein concentration was determined by Bradford assay (Thermo Scientific, 23200). Membrane proteins were denatured in Laemmli sample buffer (Thermo Scientific, J60015.AC) at 55 °C for 20 min prior to electrophoresis.

### Cycloheximide chase and inhibitor treatments

Protein turnover was measured by cycloheximide (CHX; Thermo Scientific, 357420250) chase assays. Strains were grown in GMM+YE at 37 °C for 16 h, after which CHX was added to 300 µg/ml to arrest translation. For proteasome inhibition, MG-132 (Adooq Bioscience, A11043) was added to 40 µg/ml 1 h before CHX and supplemented every 2 h at 40 µg/ml. Mycelia were collected 0, 2 and 4 h after CHX addition, unless otherwise indicated, and processed for membrane protein extraction as above. Band intensities were quantified in ImageJ and normalized to the total protein signal of the corresponding lane. Signal linearity was validated. Unless otherwise indicated in the figure legends, protein degradation was expressed as the protein level at 2 h or 4 h after CHX addition relative to the steady-level (0 h time point) of the same strain and treatment condition; experiments were performed in three biological replicates; and the relative level of Hmg1 remaining after CHX addition was compared between conditions by two-way ANOVA. Analyses were performed in GraphPad Prism (11.0.2).

### SDS-PAGE and immunoblotting

For electrophoresis, 20 µg of protein was separated by 4-15% sodium dodecyl sulfate-polyacrylamide gel (Bio-Rad) in running buffer containing 25 mM Tris, 192 mM glycine, 0.1% SDS, and transferred to nitrocellulose membranes by semi-dry transfer in 25 mM Tris, 192 mM glycine, 10% (v/v) methanol. Following transfer, membranes were stained with Ponceau S and imaged to confirm equal loading and transfer. Membranes were then destained. HiBiT-tagged proteins were detected with the Nano-Glo HiBiT Blotting System (Promega, N2410) following the manufacturer’s instructions and imaged by chemiluminescence. For antibody-based detection, membranes were blocked in 5% (w/v) non-fat milk in TTBS (50 mM Tris-HCl pH 7.4, 150 mM NaCl, 0.1% (v/v) Tween-20) and probed with specific primary antibodies, followed by incubation with horseradish peroxidase-conjugated secondary antibodies. Immunoblots were developed with Clarity Western ECL substrate (Bio-Rad, 1705060). FLAG-tagged proteins were detected with an anti-FLAG monoclonal antibody (clone FG4R; Invitrogen, MA1-91878; 1:1,000) and an HRP-conjugated goat anti-mouse secondary antibody (Invitrogen, 31430; 1:10,000). GFP-tagged proteins were detected with an anti-GFP polyclonal antibody (GenScript, A01704; 1:1,000) and an HRP-conjugated goat anti-rabbit secondary antibody (Invitrogen, 31466; 1:10,000).

### Co-immunoprecipitation

For co-immunoprecipitation, membrane fractions were prepared as described above. To pull down Hmg1 as bait, 20 µl Protein G magnetic beads (Cytiva, 28951379) were incubated with 2 µg of anti-HiBiT monoclonal antibody (Promega, N7200), and the antibody was then crosslinked to the beads with dimethyl pimelimidate (DMP) at 5.4 mg/ml in 0.2 M triethanolamine for 45 min at room temperature. Unreacted DMP was quenched with 50 mM ethanolamine for 15 min at room temperature. The crosslinked beads were incubated with 2 mg of membrane protein for 5.5 h at 4 °C with end-over-end rotation. For the pull-down with InsA as bait, anti-DYKDDDDK magnetic agarose (Invitrogen, A36797) was used according to the manufacturer’s instructions. Beads were collected on a magnetic stand, washed three times with PBS, and bound proteins were eluted in non-reducing sample buffer (120mM Tris-HCl pH 6.8, 4% SDS, 20% glycerol, 0.02% bromophenol blue) at 55 °C for 20 min followed by denaturation with 5% 2-mercaptoethanol at 55 °C for 10 min. The samples were analyzed by immunoblotting as above.

### RNA extraction and RT-qPCR analysis

For RNA extraction, mycelia were grown in liquid GMM at 37 °C and 250 r.p.m. for 16 h. Mycelia were collected and ground to a powder under liquid nitrogen. Total RNA was extracted with the RNeasy Mini Kit (Qiagen, 74106), and cDNA was synthesized using the PrimeScript FAST RT reagent Kit with gDNA Eraser (Takara Bio, RR092A). RT-qPCR was run with TB Green Premix Ex Taq II Fast qPCR (Takara Bio, RR830A) on a CFX Connect Real-Time System (Bio-Rad). Relative expression was determined by the 2^^-ΔΔCt^ method using *tubA* as the reference gene. Data are from three biological replicates, and significance was assessed by paired *t*-test on ΔCt values.

### Antifungal susceptibility testing

Voriconazole minimum inhibitory concentrations (MICs) were determined by broth microdilution according to CLSI standard M38. Conidia were suspended in RPMI 1640 medium (buffered to pH 7.0 with 0.165 M MOPS) at a final inoculum of 5 × 10[ conidia/ml. Plates were incubated at 37 °C, and MICs were read visually at 48 h as the lowest drug concentration producing complete inhibition of growth. For targeted degradation of Hmg1 during susceptibility testing, doxycycline was supplied either at the start of the assay and maintained until the endpoint reading, or after 16 h of growth and maintained until reading, as indicated for each experiment.

---

**Supplementary Figure 1.**
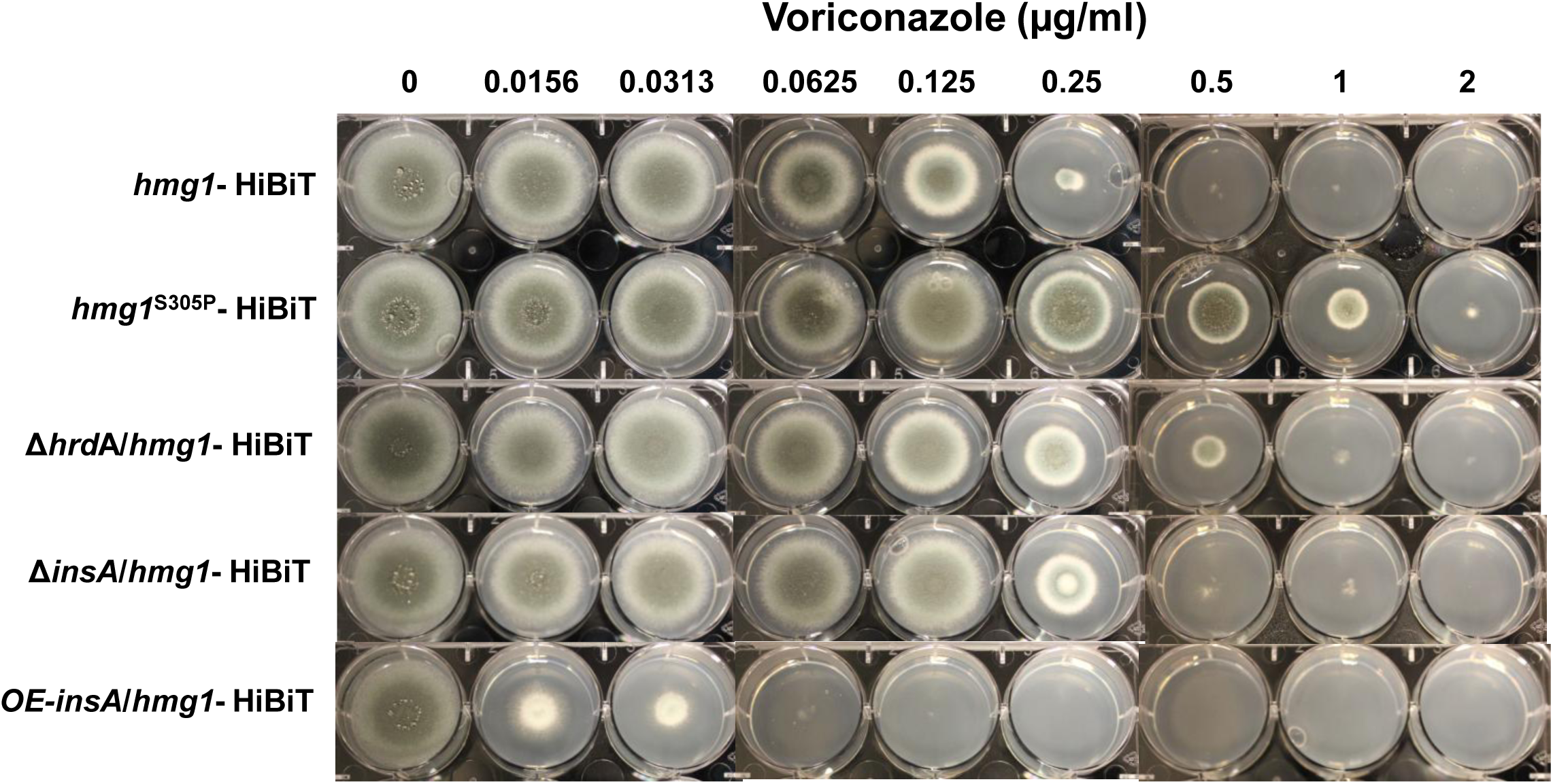
Voriconazole susceptibility profiles of strains used in this study. Radial growth assays of the indicated strains on GMM agar containing voriconazole at the concentrations shown. Conidia were spot-inoculated and plates were incubated at 37°C for 3 days.

**Supplementary Figure 2.**
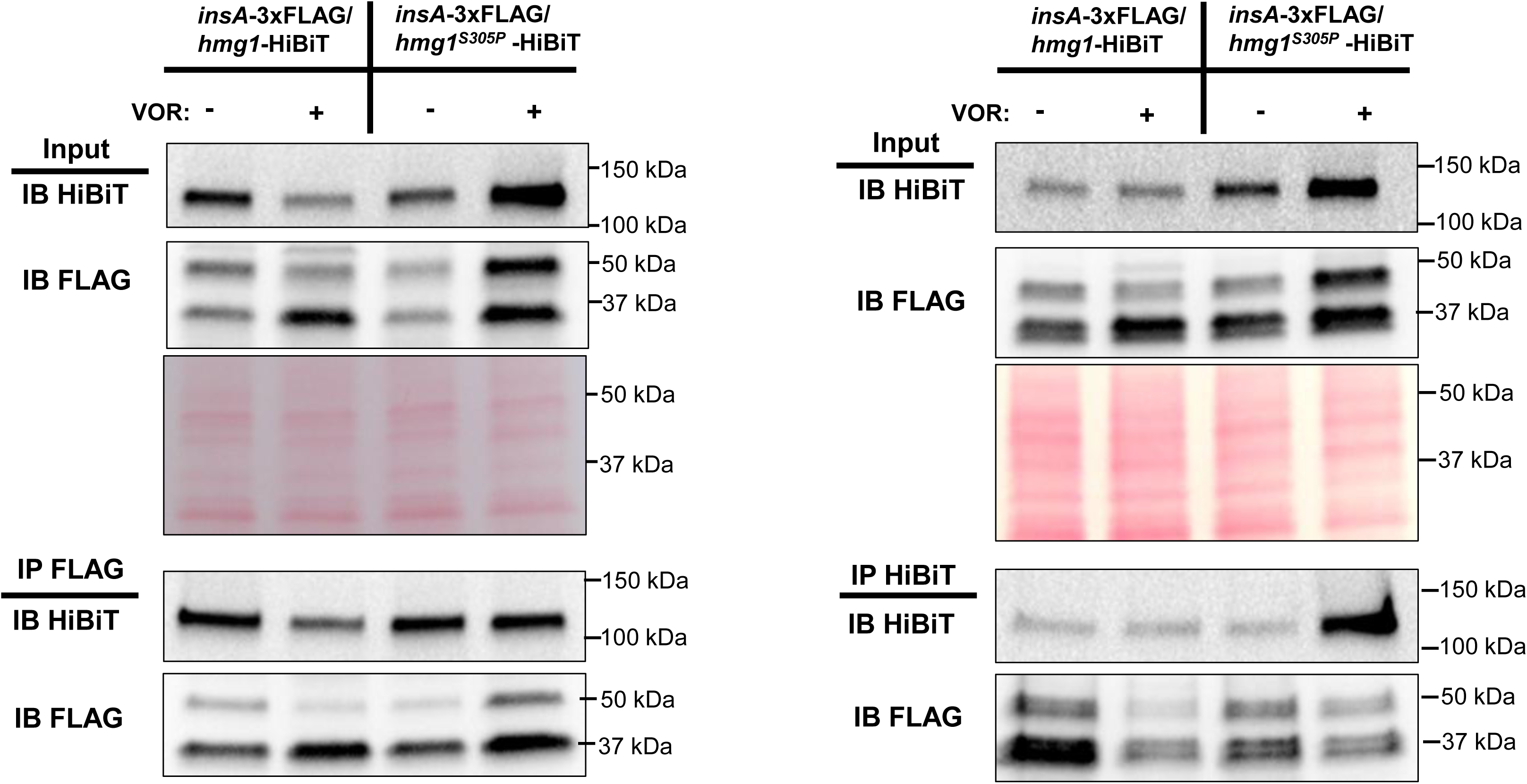
InsA remains associated with Hmg1 regardless of SSD mutation or voriconazole treatment. Co-immunoprecipitation of InsA and Hmg1 in the *insA*-3×FLAG/*hmg1*-HiBiT and *insA*-3×FLAG/*hmg1^S305P^*-HiBiT strains. Strains were grown for 16 h with or without voriconazole (VOR, 0.0625 µg/ml). Membrane fractions were prepared and subjected to immunoprecipitation with anti-FLAG magnetic agarose (left panels, IP FLAG) or anti-HiBiT antibody crosslinked to Protein G magnetic beads (right panels, IP HiBiT). Input and precipitated fractions were analyzed by immunoblotting with anti-FLAG antibodies and Nano-Glo HiBiT Blotting System.

**Supplementary Figure 3.**
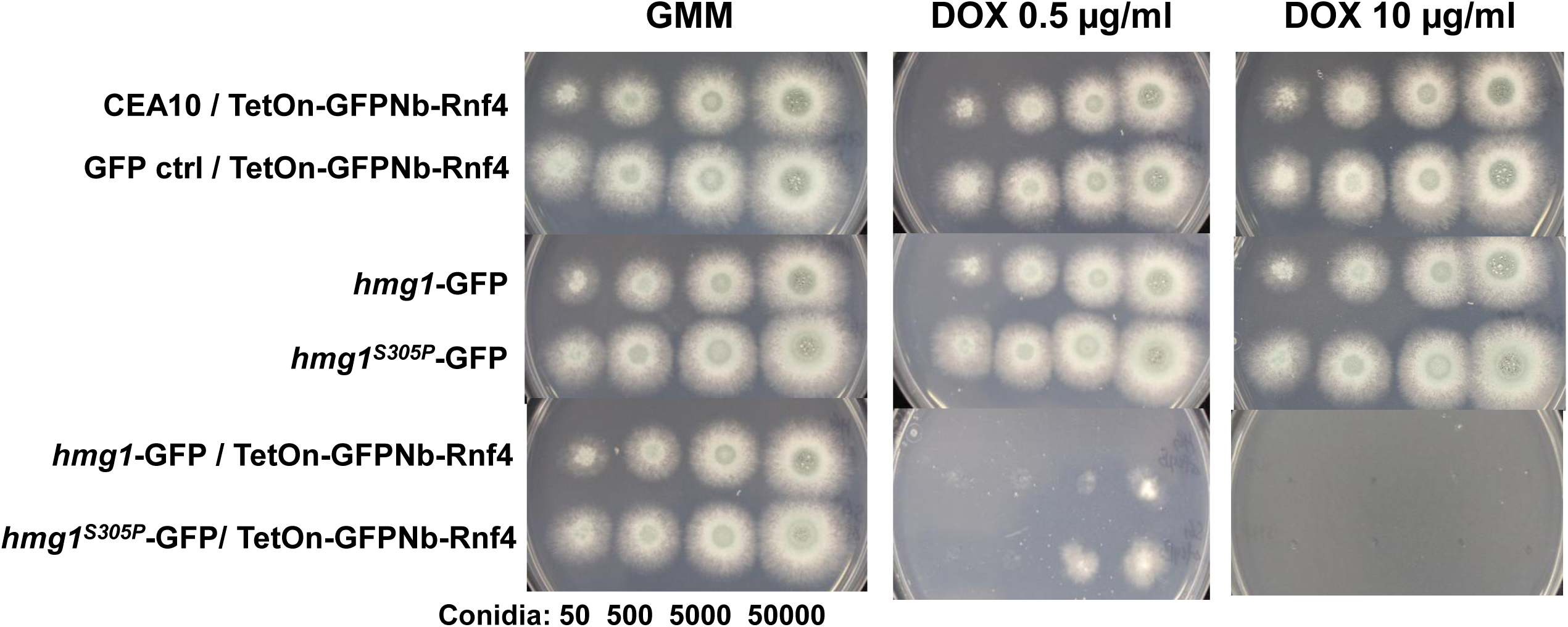
Targeted degradation of Hmg1 specifically inhibits growth. Radial growth assays of the parental strain and GFP-targeted strains. Conidia were spot-inoculated onto GMM agar containing no doxycycline, 0.5 µg/ml or 10 µg/ml doxycycline and incubated at 37°C for 2 days.

## ACKNOWLEDGEMENTS

This work was supported in part by NIH/NIAID grant R01AI195581 to JRF.

